# Degradation of cyclin B is critical for nuclear division in *Trypanosoma brucei*

**DOI:** 10.1101/213652

**Authors:** Hanako Hayashi, Bungo Akiyoshi

## Abstract

Kinetoplastids have a nucleus that contains the nuclear genome and a kinetoplast that contains the mitochondrial genome. These single-copy organelles must be duplicated and segregated faithfully to daughter cells at each cell division. In *Trypanosoma brucei*, although duplication of both organelles starts around the same time, segregation of the kinetoplast precedes that of the nucleus. Cytokinesis subsequently takes place so that daughter cells inherit a single copy of each organelle. Very little is known about the molecular mechanism that governs the timing of these events. Furthermore, it is thought that *T. brucei* lacks a spindle checkpoint that delays the onset of nuclear division in response to spindle damage. Here we show that a mitotic cyclin CYC6 has a dynamic localization pattern during the cell cycle, including kinetochore localization from G2 to metaphase. Using CYC6 as a molecular cell cycle marker, we confirmed that *T. brucei* cannot delay the onset of anaphase in response to a bipolar spindle assembly defect. Interestingly, expression of a stabilized form of CYC6 caused the nucleus to arrest in a metaphase-like state without preventing cytokinesis. We propose that trypanosomes have an ability to regulate the timing of nuclear division by modulating the CYC6 protein level, without a spindle checkpoint.

## Introduction

Accurate transmission of genetic material to offspring is essential for the survival of organisms. The genome in eukaryotes exists in different organelles such as the nucleus, mitochondria, and plastids. Nuclear DNA is duplicated during S phase and segregated equally to daughter cells during M phase. Kinetochores are the macromolecular protein complexes that assemble onto centromeric DNA and interact with spindle microtubules. It is essential that sister kinetochores attach to spindle microtubules emanating from opposite poles in metaphase so that sister chromatids segregate away from each other during anaphase. Cells possess a surveillance mechanism, called the spindle checkpoint, that delays the onset of anaphase in response to defects in kinetochore-microtubule attachments (London and Biggins 2014; Musacchio 2015). Once all sister kinetochores have achieved proper bi-oriented attachments, the spindle checkpoint is satisfied. This results in the ubiquitylation of two key targets cyclin B and securin by the anaphase-promoting complex (APC/C), leading to their destruction by proteasomes.

In contrast to nuclear DNA, the mechanism of mitochondrial DNA transmission varies among eukaryotes. For example, in animals that have a high copy number of mitochondria, transmission of mitochondrial DNA is thought to occur randomly (Westermann 2010). On the other hand, a single mitochondrion is present in many unicellular eukaryotes, such as kinetoplastids, *Plasmodium falciparum* and *Cyanidioschyzon merolae* (Robinson and Gull 1991; Itoh et al. 1997; Okamoto et al. 2009). The timing of duplication and partition of their mitochondria must be coordinated with the cell cycle machinery in these organisms. Kinetoplastids are a group of unicellular organisms that are characterized by the unique structure called the kinetoplast, which is a network of multiple copies of mitochondrial DNA (termed the kDNA) enclosed in a single mitochondrion (Vickerman 1962). They are evolutionarily divergent from commonly studied model eukaryotes (e.g. yeast, worms, flies, and humans) (Cavalier-Smith 2010; Walker et al. 2011), so understanding their biology can provide insights into the extent of conservation or divergence in eukaryotes. Among various kinetoplastids studied thus far, the mechanism of cell cycle is best characterized in *Trypanosoma brucei*, the causative agent of human African trypanosomiasis (for reviews, see (McKean 2003; Hammarton 2007; Vaughan and Gull 2008; Li 2012)). *T. brucei* has a canonical cell cycle for nuclear events (G1, S, G2, and M phases). G1 cells have a single kinetoplast and nucleus (termed 1K1N). Duplication of kinetoplast DNA starts almost simultaneously with that of nuclear DNA, but completes earlier (Woodward and Gull 1990; Siegel et al. 2008). Segregation of kDNA depends on that of basal bodies and occurs during the nuclear S phase, creating 2K1N cells (Robinson and Gull 1991; Ogbadoyi et al. 2003). Trypanosomes do not break down their nuclear envelope (closed mitosis), and an intranuclear mitotic spindle is assembled in the nucleus during M phase (Vickerman and Preston 1970; Ogbadoyi et al. 2000). Sister kinetochores align at the metaphase plate during metaphase, followed by the separation of nuclear DNA in anaphase (creating 2K2N cells) and split of cells by cytokinesis (Sherwin and Gull 1989; Woodward and Gull 1990). It is essential that replication and segregation of these organelles occur prior to cytokinesis in a coordinated manner so that daughter cells inherit a copy of each. Little is known about the underlying molecular mechanism.

Available evidence suggests that *T. brucei* is not capable of halting their cell cycle in response to various defects in the nucleus. For example, when bipolar spindle assembly is blocked in procyclic (insect form) cells, they undergo cytokinesis without a noticeable delay despite a lack of nuclear division (Robinson et al. 1995; Ploubidou et al. 1999). This results in the formation of one daughter cell that has one kinetoplast DNA without nuclear DNA (1K0N, termed zoid) and another cell that has one kinetoplast with tetraploid DNA content, suggesting that the spindle checkpoint is not operational (Ploubidou et al. 1999). In fact, most of the spindle checkpoint components (i.e. Mps1, Mad1, Mad3/BubR1, Bub1, Bub3) are not found in *T. brucei* or other kinetoplastids. Although a Mad2 homolog is present, this protein localizes at basal bodies, not kinetochores (Akiyoshi and Gull 2013). It is therefore thought that trypanosomes cannot delay cytokinesis even when nuclear division fails to occur. Yet, there must be a mechanism to coordinate the segregation of nuclear DNA with cytokinesis in unperturbed cells. One possibility is the presence of a cell cycle oscillator that triggers cell cycle events in a set sequence even without feedback control systems. The best characterized components of cell cycle oscillators are cyclin/CDK (cyclin-dependent kinase) complexes (Nurse 1990; Morgan 1997; Gérard et al. 2015). The rise and fall of their kinase activities trigger cell cycle events in a set sequence. For example, increased activities of mitotic CDK complexes promote entry into M phase and various mitotic events, whereas their decrease is essential for exit from mitosis. *T. brucei* has ten cyclins and eleven CDKs, among which CYC6/CRK3 is the major mitotic cyclin/CDK complex in *T. brucei* (CYC6 is also known as CycB2) (Li and Wang 2003; Hammarton et al. 2003). When degradation of CYC6 was inhibited by proteasome inhibitors or APC/C downregulation, cells accumulated in a metaphase-like state with a bipolar spindle (Mutomba et al. 1997; Kumar and Wang 2005). These observations suggested that degradation of cyclin B could be a trigger for the metaphase-anaphase transition. Here we directly tested this possibility by expressing a non-degradable version of CYC6 in *T. brucei*.

## Results

### Identification of cyclin B^CYC6^ as a molecular cell cycle marker

Cellular localization of CYC6 has not been reported thus far, so we first examined it by endogenously tagging CYC6 with an N-terminal YFP tag in *T. brucei* procyclic cells. We observed the following localization pattern (Figure 1A). There was no distinct signal in G1 cells. From S phase onwards, CYC6 was found at the basal body area and flagellum. From G2 to metaphase, nuclear signal was observed with significant enrichment at kinetochore regions in metaphase. In fact, these nuclear dots co-localized with a kinetochore marker protein, KKT2 (Figure 1B). CYC6 disappeared from the nucleus in anaphase. We obtained similar results for CRK3, which formed dots in metaphase and disappeared in anaphase (Figure 1C). Thus, CYC6 and CRK3 exhibit a differential localization pattern depending on cell cycle stages, and can therefore be used as a molecular cell cycle marker.

**Figure 1.**
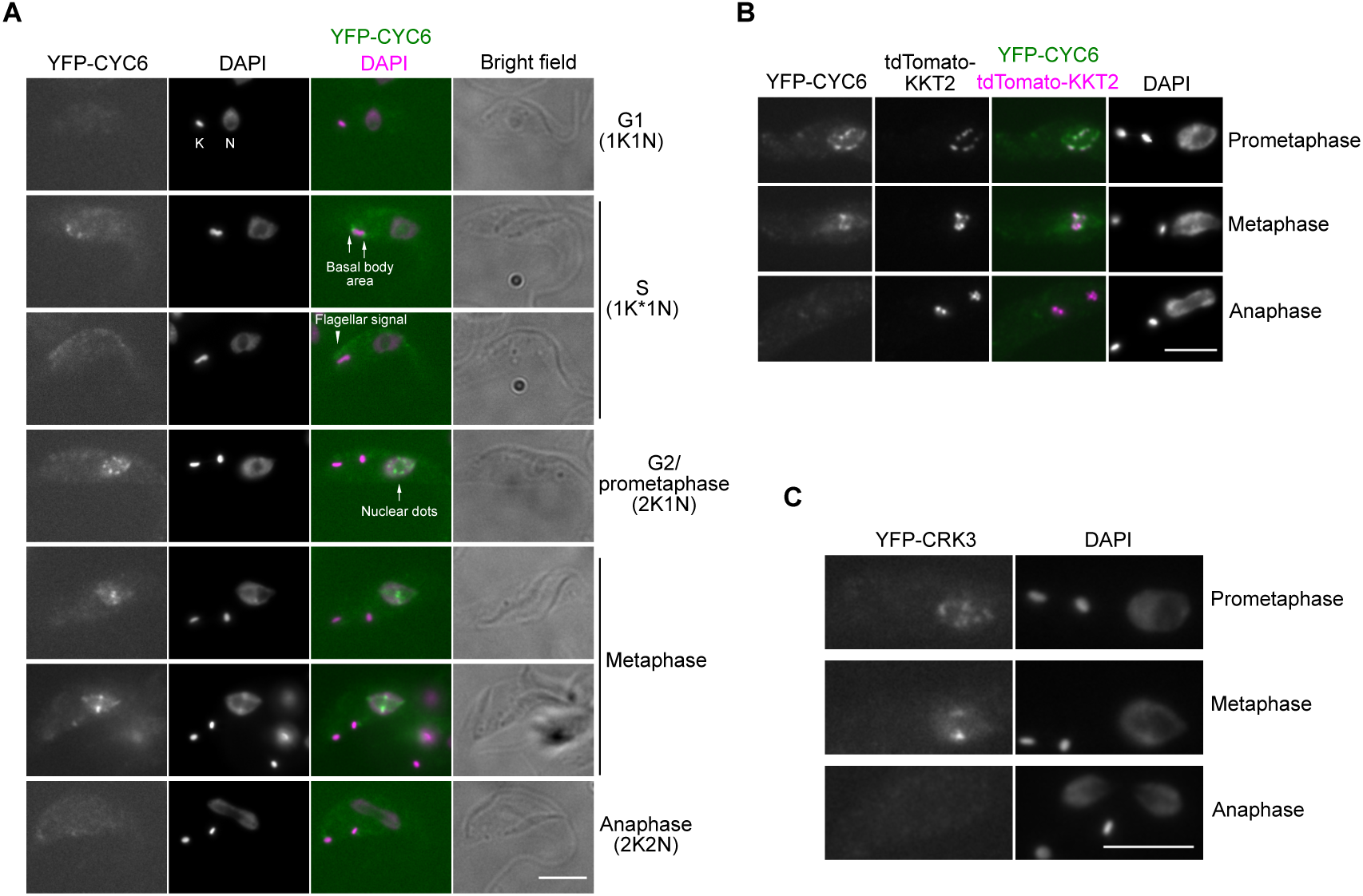
Cyclin B^CYC6^ is enriched at kinetochores in metaphase and disappears in anaphase. (A) CYC6 has a dynamic localization pattern during the cell cycle. Examples of procyclic form cells that express YFP-CYC6 are shown (cell line BAP426). K and N stands for the kinetoplast and nucleus, respectively. (B) CYC6 nuclear dots partially co-localize with a kinetochore protein, KKT2 (BAP1005). (C) CRK3 has nuclear dots in metaphase and disappears in anaphase (BAP463). Bars, 5 µm.

### Cyclin B^CYC6^ is important for bipolar spindle assembly, but not for kinetochore assembly

CDK activities are known to be important for kinetochore assembly in some eukaryotes, including humans (Gascoigne and Cheeseman 2013). The finding that CYC6 localizes at kinetochores from G2 to metaphase in trypanosomes prompted us to study its importance for kinetochore assembly. We therefore depleted CYC6 by RNAi-mediated knockdown (Ngô et al. 1998). We confirmed that CYC6 is essential for cell growth, as previously reported (Li and Wang 2003; Hammarton et al. 2003) (data not shown). Because cyclin/CDK activities are known to be important for various mitotic events (Bishop et al. 2000), we first examined bipolar spindle formation. We used a spindle marker protein that we identified from our previous tagging screen (ORF Tb927.11.14370) (Archer et al. 2011; Akiyoshi and Gull 2014). This protein had a localization pattern characteristic of spindle microtubules, so we named it MAP103 for <u>m</u>icrotubule-<u>a</u>ssociated <u>p</u>rotein <u>103</u> kDa (Figure S1). We observed defective spindle microtubules in CYC6-depleted cells, suggesting that CDK activities are essential for proper bipolar spindle assembly (Figure 2A). Under these conditions, however, localization of all KKT proteins we examined was not affected (KKT1, KKT4, KKT7, KKT8, KKT10, KKT14, KKT16) (Figure 2B). Therefore, CYC6 is dispensable for the assembly of these kinetochore proteins in procyclic cells.

**Figure 2.**
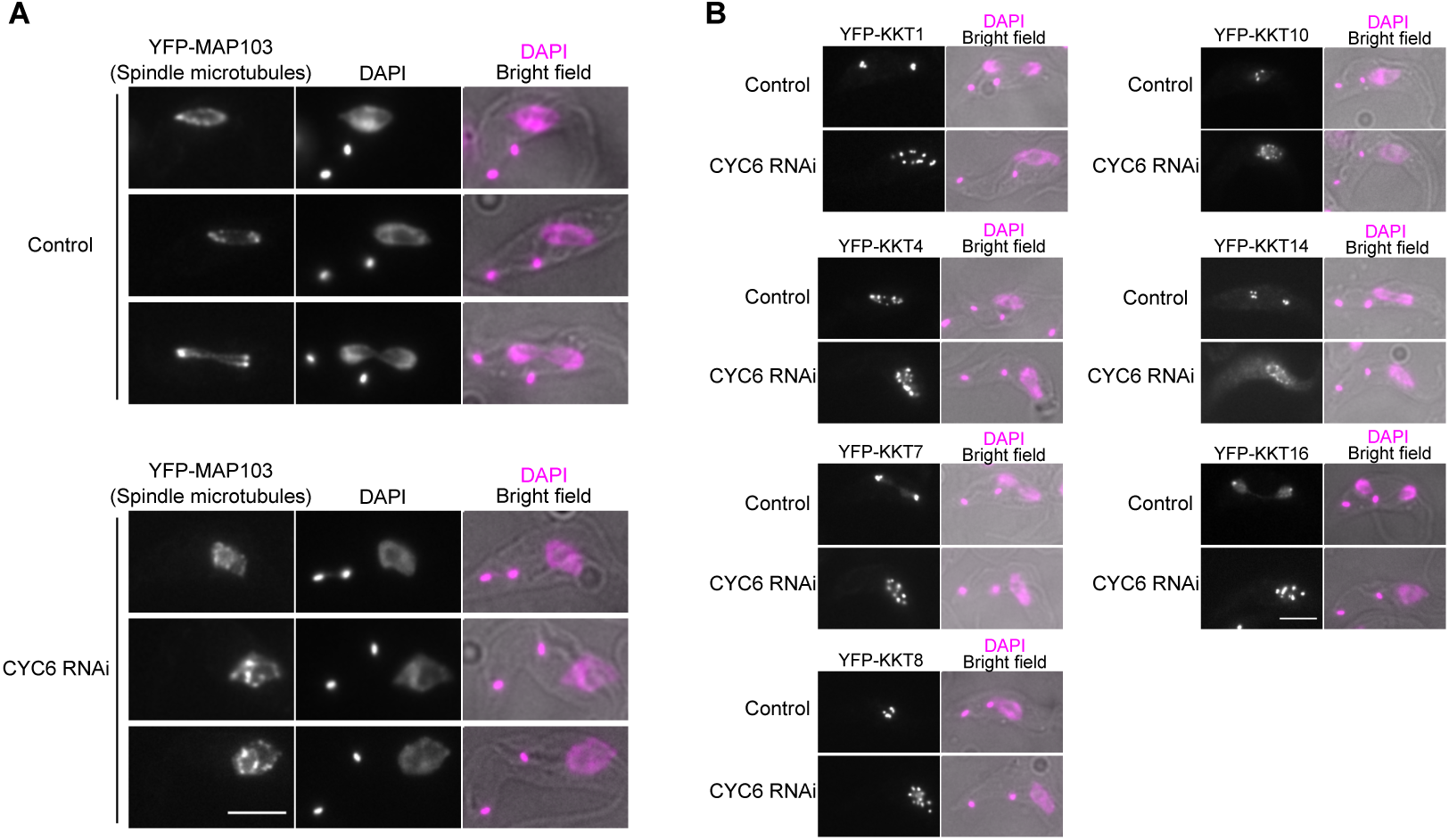
Cyclin B^CYC6^ is important for bipolar spindle assembly, but dispensable for kinetochore assembly. (A) Bipolar spindle formation was perturbed upon induction of CYC6 RNAi. Cells expressing YFP-MAP103 (a marker for spindle microtubules) were fixed at 24 hr post-induction (BAP504). (B) Kinetochore localization of KKT1, KKT4, KKT7, KKT8, KKT10, KKT14, and KKT16 proteins was not affected by CYC6 depletion (BAP503, BAP585, BAP505, BAP593, BAP596, BAP506, and BAP604, respectively). Examples of 2K1N (prometaphase/metaphase) or 2K2N (anaphase) cells expressing indicated YFP-KKT proteins fixed at 24 hr post-induction are shown. Bars, 5 µm.

### Cells fail to delay the onset of anaphase in response to spindle defects

We next used CYC6 as a molecular cell cycle marker to examine the effect of drugs. We first used an anti-microtubule agent, ansamitocin, to examine the effect of a bipolar spindle assembly defect for cell cycle progression (Robinson and Gull 1991). By testing various concentrations of ansamitocin, we found that 5 nM of ansamitocin significantly slowed down cell growth (Figure 3A). After a 4-hr treatment, nuclear division and bipolar spindle assembly was perturbed as expected (Figure 3B). In this condition, however, we found no significant enrichment of nuclear CYC6-positive cells (Figure 3C). This corroborates previous studies (Ploubidou et al. 1999) and confirms that trypanosomes are not capable of delaying the onset of anaphase in response to spindle damage.

**Figure 3.**
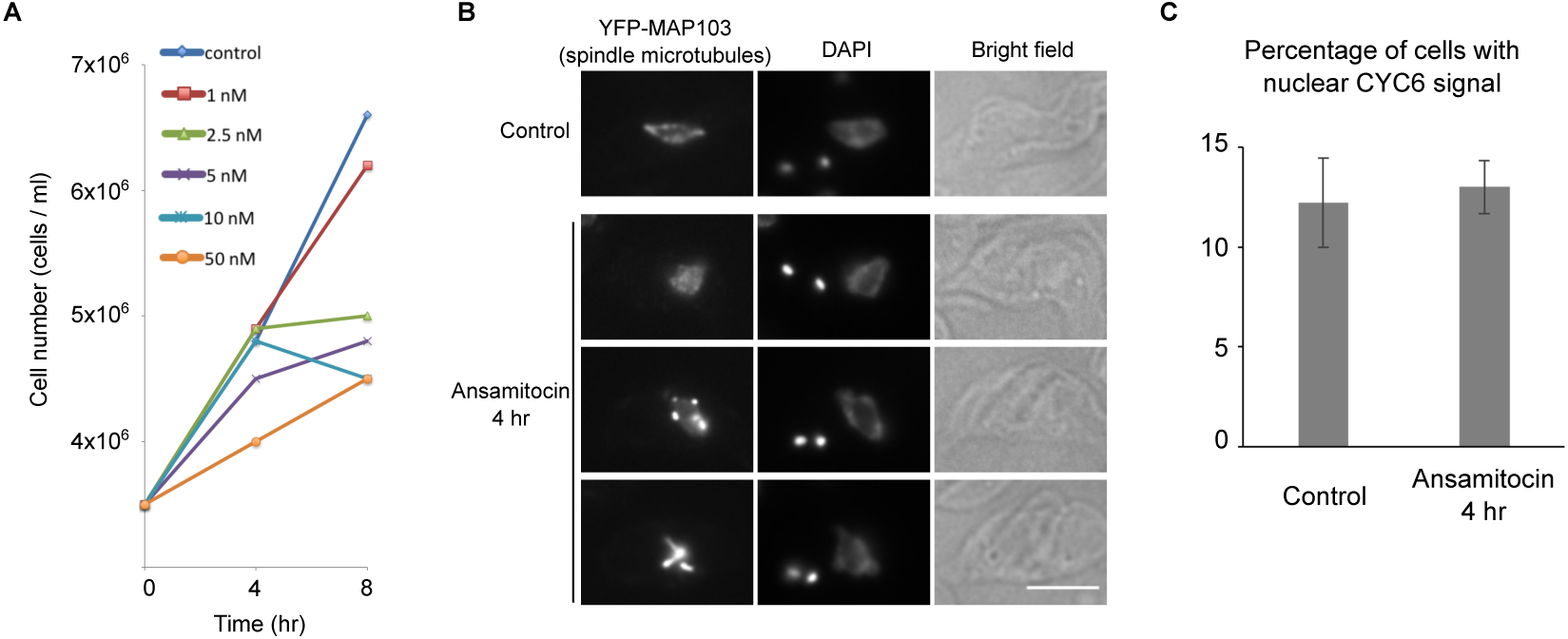
Spindle assembly defects do not cause cyclin B^CYC6^ accumulation in the nucleus. (A) Growth curves of control and ansamitocin-treated cultures show a concentration-dependent growth inhibition (BAP125). (B) Ansamitocin prevents bipolar spindle assembly. Cells expressing YFP-MAP103 (BAP79) were treated with 5 nM ansamitocin for 4 hr and fixed. Bar, 5 µm. (C) Ansamitocin treatment does not result in the accumulation of nuclear CYC6-positive cells. Cells expressing YFP-CYC6 (BAP426) were treated with 5 nM ansamitocin for 4 hr and fixed. Three hundred cells were counted for each sample, and experiments were performed three times. Error bars represent standard deviation.

### Stabilization of cyclin B^CYC6^ causes metaphase arrest in the nucleus

We next examined the effect of cyclin B stabilization for cell cycle progression. We first used a proteasome inhibitor MG-132 that blocked cell cycle progression and stabilized the CYC6 protein (Mutomba et al. 1997; Bessat et al. 2013). When cells expressing YFP-CYC6 were treated with 10 µM MG-132 for 4 hr, ~30% of cells had nuclear CYC6 signal (compared to ~10 % in control), suggesting that the nucleus arrested prior to anaphase (Figure 4A, B). Indeed, these cells had a bipolar spindle (often elongated) and most of their kinetochores were aligned at the metaphase plate (Figure 4C, D). We also noted that the distance between the two kinetoplast DNA in these cells was often greater than that in control metaphase cells. These results suggest that, upon MG-132 treatment, trypanosomes arrest the nucleus in a metaphase-like state in which cyclin B is not degraded, although their cytoplasm transits to an anaphase-like state.

**Figure 4.**
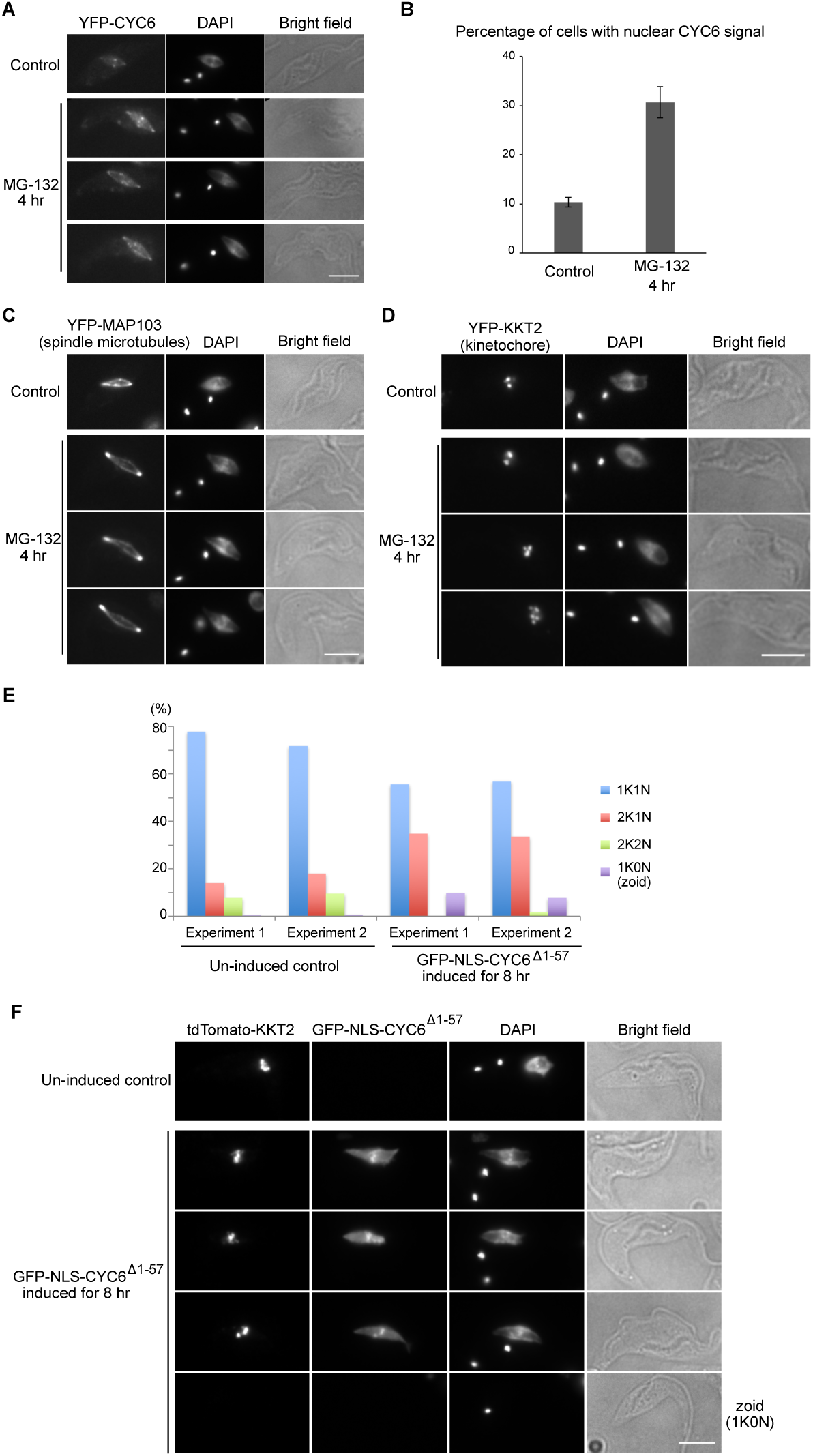
Cyclin B^CYC6^ prevents nuclear division. (A–D) MG-132 treatment causes metaphase arrest. Cells expressing YFP-CYC6 (A, B: BAP426), YFP-MAP103 (C: BAP79), or YFP-KKT2 (D: BAP122) were treated with 10 µM MG-132 for 4 hr and fixed, showing that a higher ratio of cells have nuclear CYC6 signal with a bipolar spindle and aligned kinetochores upon MG-132 treatment. For quantification of nuclear CYC6-positive cells (B), 300 cells were counted for each sample, and experiments were performed three times. Error bars represent standard deviation. (E, F) Expression of a non-degradable CYC6 protein in the nucleus delays nuclear division. GFP-NLS-CYC6^Δ1–57^ expression was induced with 0.1 µg/ml doxycycline in cells that have tdTomato-KKT2 (BAP945) for 8 hr. Four hundred cells were counted for each sample, and experiments were performed twice. Bars, 5 µm.

Because MG-132 treatment affects the protein level of many other proteins, we next tested whether the presence of cyclin B in the nucleus is sufficient to prevent nuclear division. Overexpression of wild-type CYC6 did not affect cell growth (data not shown). We therefore expressed a GFP-NLS fusion of a non-degradable form of CYC6 (CYC6^Δ1–57^). Interestingly, we detected a decrease in 2K2N cells and accumulation of 2K1N cells upon expression of non-degradable CYC6 for 8 hr (Figure 4E), suggesting that the nucleus was arrested in a metaphase-like state. Indeed, kinetochores were aligned at the metaphase plate in these cells (Figure 4F). We also detected a significant increase in the number of zoids (1K0N cells). This implies that cytokinesis occurred despite the lack of nuclear division (Figure 4E, F). These results show that CYC6 is capable of arresting the nucleus in a metaphase-like state, although it cannot stop cytokinesis. Taken together, our data show that trypanosomes have an ability to control the timing of nuclear division by modulating the degradation of a mitotic cyclin in the nucleus.

## Discussion

Previous studies observed the formation of zoids despite a lack of nuclear division due to spindle damage (Ploubidou et al. 1999), cyclin/CDK depletion (Hammarton et al. 2003; Li and Wang 2003; Tu and Wang 2004), or expression of a non-degradable cohesin subunit SCC1 (Gluenz et al. 2008). These studies strongly suggested that *T. brucei* cannot prevent cytokinesis in response to a lack of nuclear division at least in procyclic cells (although this is likely to be the case in bloodstream form too, see (Gluenz et al. 2008)). In this study, we established CYC6 as a molecular marker for cell cycle progression, and confirmed that trypanosomes indeed failed to delay the anaphase onset in response to spindle damage. This implies that the timing mechanism of the nuclear cell cycle progression is likely governed by an intrinsic cell cycle timer, as observed in embryonic divisions (Yang and Ferrell 2013; Yuan and O’Farrell 2015) and in spindle checkpoint mutants of yeasts and flies (Hoyt et al. 1991; Li and Murray 1991; Buffin et al. 2007).

Interestingly, we found that expression of non-degradable cyclin B can delay the onset of anaphase (in the nucleus). This means that trypanosomes could potentially coordinate the timing of nuclear division with that of cytokinesis by regulating the level of the CYC6 protein in the nucleus. Because APC/C is responsible for the degradation of mitotic cyclins, understanding its regulatory mechanism is of critical importance. It is interesting to note that two kinetochore proteins (KKT4 and KKT20) co-purified with several components of the APC/C (Akiyoshi and Gull 2014; Nerusheva and Akiyoshi 2016), suggesting that kinetochores may directly regulate APC/C activities. It will be important to understand the underlying mechanism.

It remains unclear how the timing of cytokinesis onset is determined in trypanosomes. It has been suggested that it may be the segregation of basal bodies, rather than that of the nucleus, that is linked to cytokinesis in trypanosomes (Ploubidou et al. 1999). Interestingly, CYC6 signal was found not only at kinetochores but also at basal bodies and flagella. Therefore, CYC6 might also have an ability to regulate the onset of cytokinesis, which will need to be tested in future studies.

## Supplemental material

Supplemental material contains Figure S1, and Tables S1, S2, and S3.

## Materials and methods

### Trypanosome cells

All trypanosome cell lines used in this study were derived from *T. brucei* SmOxP927 procyclic form cells (TREU 927/4 expressing T7 RNA polymerase and the tetracycline repressor to allow inducible expression) (Poon et al. 2012) and are listed in Table S1. Cells were grown at 28 ºC in SDM-79 medium supplemented with 10% (v/v) heat-inactivated fetal calf serum (Brun and Schönenberger 1979). Cell growth was monitored using a CASY cell counter and analyzer system (Roche). RNAi was induced with doxycycline at a final concentration of 1 µg/ml. Non-degradable CYC6 was expressed with doxycycline at 0.1 µg/ml. Ansamitocin P-3 was purchased from Abcam (catalog number, ab144546) and MG-132 was purchased from Merck (catalog number, 474790).

### Tagging, cloning, transfections, and microscopy

Plasmids and primers used in this study are listed in Table S2 and S3, respectively. Endogenous tdTomato tagging was performed using pBA148 (Akiyoshi and Gull 2014). YFP tagging was performed using pEnT5-Y (for KKTs and MAP103) or pBA106 (for CYC6 and CRK3) tagging vectors. pBA106 is a modified version of the pEnT5-Y vector (Kelly et al. 2007) to allow N-terminal 3FLAG-6HIS-YFP tagging. A targeting sequence for the CRK3 tagging (consisting of *Xba*I site, 4–250 bp of the CRK3 coding sequence, *Not*I site, 250 bp of CRK3 5’UTR, *Bam*HI site) was synthesized by GeneArt. To make pBA106, a synthetic DNA fragment that encodes a 3FLAG-6HIS tag (made by annealing BA403 and BA404) was ligated into pEnT5-Y using *Hind*III and *Spe*I sites. For generation of the inducible CYC6 RNAi cell line, 424 bp fragment targeting 378–801 bp of the CYC6 coding sequence was amplified from genomic DNA and cloned into the p2T7-177 vector (Wickstead et al. 2002), creating pBA734. To make a non-degradable version of CYC6 with an N-terminal GFP-NLS tag (pBA1319: GFP-NLS-CYC6^Δ1–57^), DNA fragment encoding CYC6^58–426^ was amplified from genomic DNA and cloned into pBA310 (Nerusheva and Akiyoshi 2016) using *Pac*I and *Asc*I sites. Plasmids linearized by *Not*I were transfected to trypanosomes by electroporation into an endogenous locus (pEnT5-Y, pBA106, and pBA148 derivatives) or 177 bp repeats on minichromosomes (p2T7-177 and pBA310 derivatives). Transfected cells were selected by the addition of 25 µg/ml hygromycin (pEnT5-Y and pBA106 derivatives), 10 µg/ml blasticidin (pBA148 derivatives), or 5 µg/ml phleomycin (p2T7-177 and pBA310 derivatives). Microscopy was performed essentially as previously described using a Leica DM5500 B microscope (Leica Microsystems) housed in the Keith Gull’s laboratory (Akiyoshi and Gull 2014) to image YFP-MAP103 or DeltaVision fluorescence microscope (Applied Precision) housed in the Micron Oxford Advanced Bioimaging Unit (Nerusheva and Akiyoshi 2016) for all other experiments.

## Acknowledgments

We thank Keith Gull for discussion, and Bela Novak, Olga Nerusheva, Gabriele Marcianò, Midori Kanazawa, and Patryk Ludzia for comments on the manuscript. We also thank the Micron Oxford Advanced Bioimaging Unit. H.H. was supported by a Uehara Memorial Foundation fellowship. B.A. was supported by a Sir Henry Dale Fellowship jointly funded by the Wellcome Trust and the Royal Society (grant number 098403/Z/12/Z), Wellcome-Beit Prize Fellowship (grant number 098403/Z/12/A), and the EMBO Young Investigator Program.

## Author’s contributions

B.A. conceived and designed the project. H.H. performed experiments for Figure 1B and Figure 4E, 4F. B.A. performed the rest of experiments, analyzed data, and wrote the manuscript.

## Supplemental material, Hayashi and Akiyoshi

**Figure S1.**
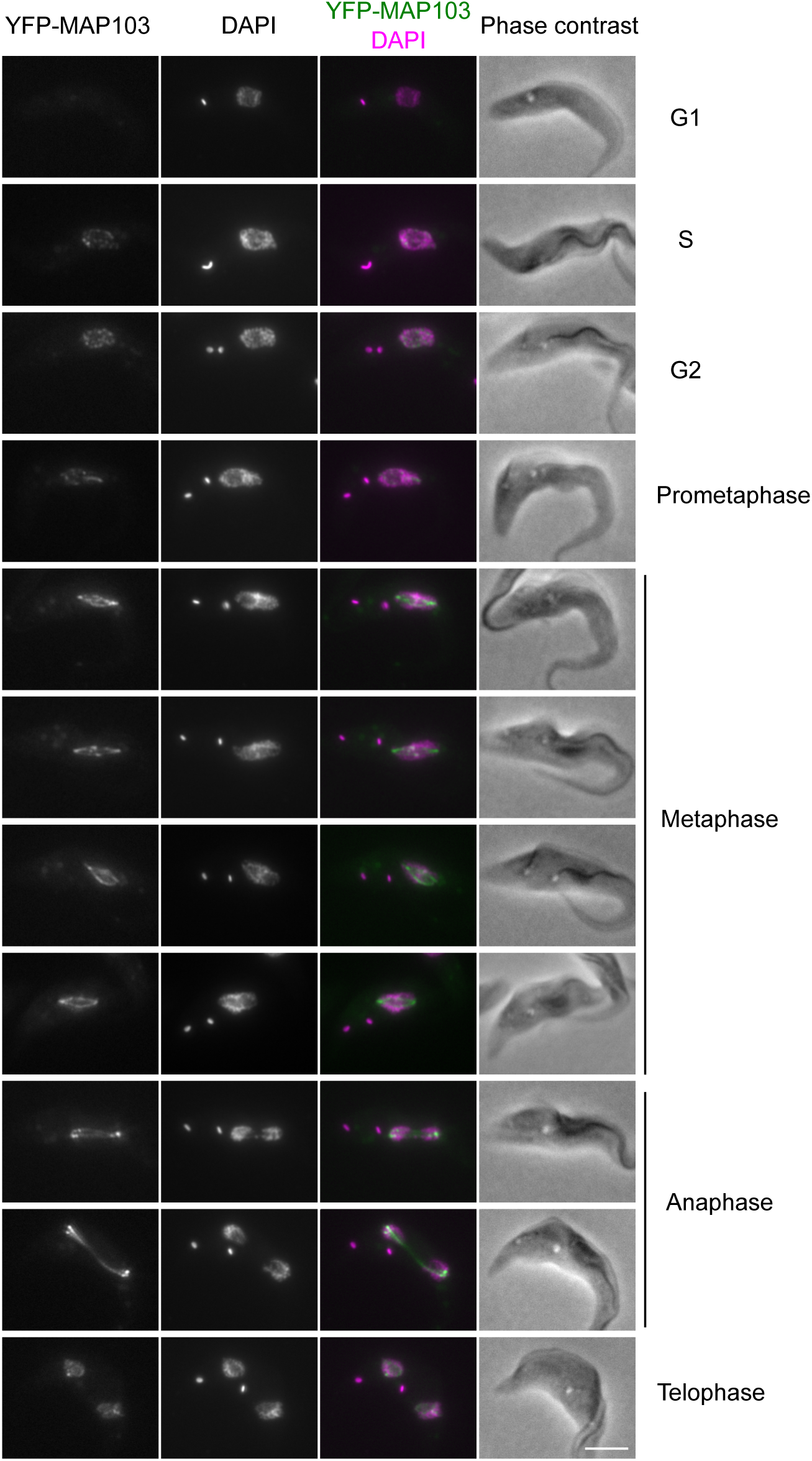
MAP103 localizes onto spindle microtubules during mitosis. Examples of cells expressing YFP-MAP103 at indicated cell cycle stages are shown. Bar, 5 µm.

**Table S1.** Trypanosome cell lines used in this study.

| Strain | Description |
| --- | --- |
| SmOxP9 | Parental cell line that expresses TetR and T7 RNAP (Kelly et al. 2007) |
| BAP79 | TY-YFP-MAP103 (this study) |
| BAP122 | TY-YFP-KKT2 (Akiyoshi and Gull 2014) |
| BAP125 | TY-YFP-KKT4 (Akiyoshi and Gull 2014) |
| BAP426 | 3FLAG-6HIS-YFP-CYC6 (this study) |
| BAP463 | 3FLAG-6HIS-YFP-CRK3 (this study) |
| BAP503 | TY-YFP-KKT1, Inducible CYC6 RNAi (this study) |
| BAP504 | TY-YFP-MAP103, Inducible CYC6 RNAi (this study) |
| BAP505 | TY-YFP-KKT7, Inducible CYC6 RNAi (this study) |
| BAP506 | TY-YFP-KKT14, Inducible CYC6 RNAi (this study) |
| BAP585 | TY-YFP-KKT4, Inducible CYC6 RNAi (this study) |
| BAP593 | TY-YFP-KKT8, Inducible CYC6 RNAi (this study) |
| BAP596 | TY-YFP-KKT10, Inducible CYC6 RNAi (this study) |
| BAP604 | TY-YFP-KKT16, Inducible CYC6 RNAi (this study) |
| BAP945 | TY-tdTomato-KKT2, Inducible GFP-NLS-CYC6<sup>Δ1-57</sup> (this study) |
| BAP1005 | 3FLAG-6HIS-YFP-CYC6, TY-tdTomato-KKT2 (this study) |

**Table S2.** Plasmids used in this study.

| Name | Descriptions |
| --- | --- |
| pEnT5-Y | TY-YFP tagging vector, Hygromycin (Kelly et al. 2007) |
| p2T7-177 | Inducible RNAi vector, integrate at 177 bp repeats (Wickstead et al. 2002) |
| pBA18 | TY-YFP-KKT1 tagging vector, Hygromycin (Akiyoshi and Gull 2014) |
| pBA31 | TY-YFP-MAP103 tagging vector, Hygromycin (this study) |
| pBA67 | TY-YFP-KKT2 tagging vector, Hygromycin (Akiyoshi and Gull 2014) |
| pBA68 | TY-YFP-KKT8 tagging vector, Hygromycin (Akiyoshi and Gull 2014) |
| pBA71 | TY-YFP-KKT4 tagging vector, Hygromycin (Akiyoshi and Gull 2014) |
| pBA72 | TY-YFP-KKT7 tagging vector, Hygromycin (Akiyoshi and Gull 2014) |
| pBA74 | TY-YFP-KKT10 tagging vector, Hygromycin (Akiyoshi and Gull 2014) |
| pBA96 | TY-YFP-KKT16 tagging vector, Hygromycin (Akiyoshi and Gull 2014) |
| pBA97 | TY-YFP-KKT14 tagging vector, Hygromycin (Akiyoshi and Gull 2014) |
| pBA106 | 3FLAG-6HIS-YFP tagging vector, Hygromycin (this study) |
| pBA148 | TY-tdTomato tagging vector, Blasticidin (Akiyoshi and Gull 2014) |
| pBA164 | TY-tdTomato-KKT2 tagging vector, Blasticidin (Nerusheva and Akiyoshi 2016) |
| pBA310 | Inducible expression vector, integrate at 177 bp, Phleomycin (Nerusheva and Akiyoshi 2016) |
| pBA586 | 3FLAG-6HIS-YFP-CYC6 tagging vector, Hygromycin (this study) |
| pBA670 | 3FLAG-6HIS-YFP-CRK3 tagging vector, Hygromycin (this study) |
| pBA734 | p2T7-177, CYC6 RNAi, integrate at 177 bp, Phleomycin (this study) |
| pBA1319 | Inducible GFP-NLS-CYC6<sup>Δ1-57</sup> expression vector, integrate at 177bp, Phleomycin (this study) |

**Table S3.**
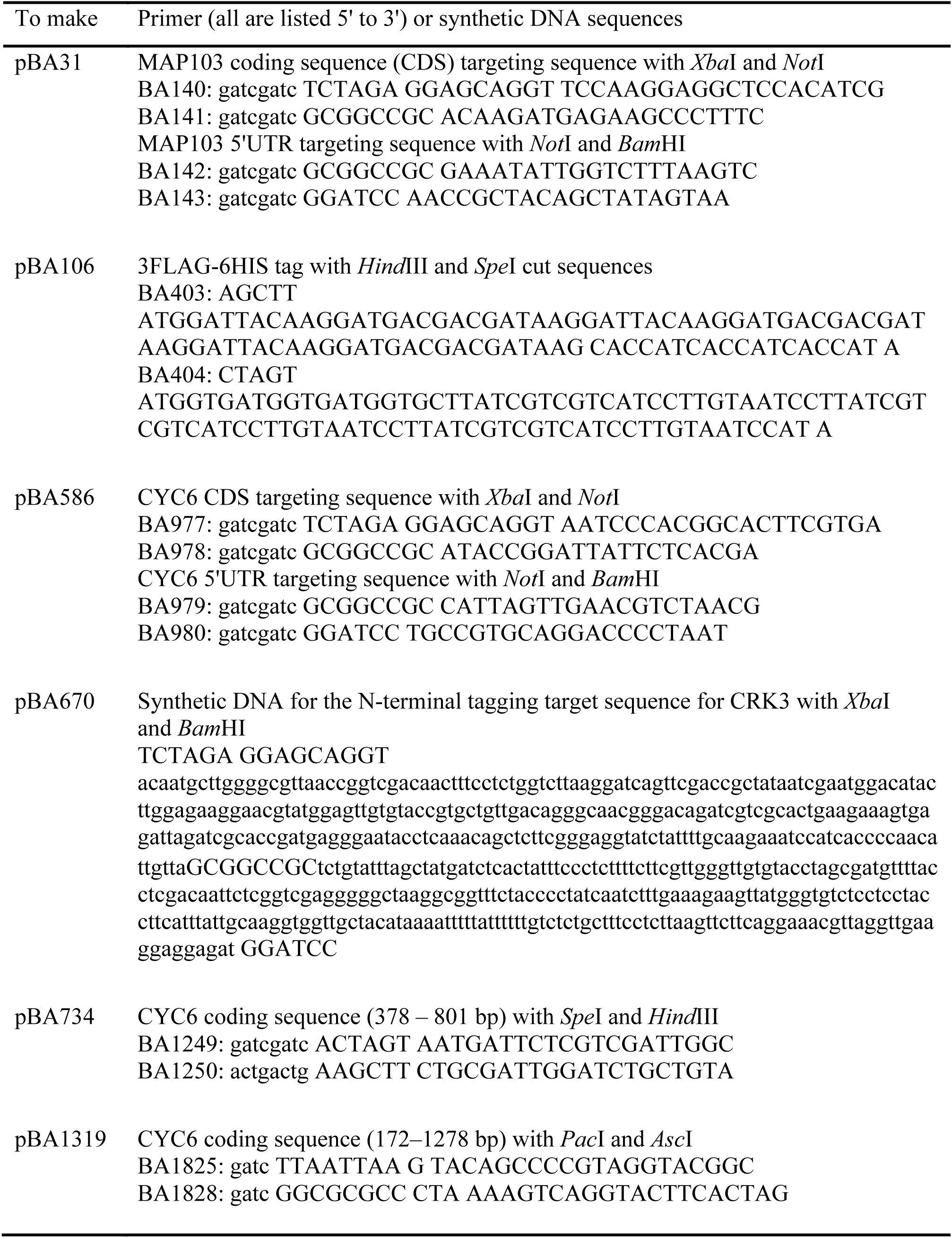
Primers and synthetic DNA sequences used in this study.

